# Mitochondrial ATP Synthase Trafficking along Microtubules to Cell Surface Depends on KIF5B and DRP1

**DOI:** 10.1101/2020.10.23.352385

**Authors:** Yi-Wen Chang, T. Tony Yang, Min-Chun Chen, Y-geh Liaw, Chieh-Fan Yin, Ting-Yu Huang, Jen-Tzu Hou, Yi-Hsuan Hung, Chia-Lang Hsu, Hsuan-Cheng Huang, Hsueh-Fen Juan

## Abstract

Ectopic adenosine triphosphate (eATP) synthase, located on the cell surface instead of the mitochondrial inner membrane, has been discovered to be a novel target for the treatment of various types of cancer. However, the mechanism of eATP synthase trafficking toward the cell surface remains unclear. In this study, we used an integrative approach incorporating multiomics, super-resolution imaging, real-time live-cell tracing, and various functional analyses to derive a transport model of eATP synthase from the mitochondria to the plasma membrane in cancer cells. We determined that the ATP synthase complex is first assembled in the mitochondria and subsequently delivered to the cell surface through the microtubule-mediated mitochondrial transport pathway. We also found that dynamin-related protein 1 (DRP1) enhances mitochondrial fission and subsequently associates with microtubule motor protein kinesin family member 5B (KIF5B) to promote the subcellular transportation of eATP synthase. Consequently, our work provides a blueprint for eATP synthase trafficking from the mitochondria toward the cell surface.

## Introduction

Adenosine triphosphate (ATP) synthase, a ubiquitous multimeric protein complex, is an important catalytic enzyme for generating ATP, the common “energy currency” of cells. This process occurs by facilitating the phosphorylation of adenosine diphosphate throughexploiting a transmembrane proton gradient (Cross, 1994). ATP synthase consists of two discrete domains: the F_O_ domain is embedded within the mitochondrial inner membrane and comprises a proton pore, and the F_1_ domain is located in the mitochondrial matrix and exhibits catalytic activity (Devenish *et al*, 2008; Jonckheere *et al*, 2012). The c-ring oligomer of the F_O_ domain rotates when protons flux through it, after which the central stalk of the ATP synthase-γ subunit drives the rotation of the F_1_ domain. The α3β3 subunits of the F_1_ domain then undergo a series of conformational changes that lead to the synthesis of ATP (Pu *et al*, 2008).

In previous studies, ATP synthase was generally found to be embedded in the inner membrane of mitochondria. Nevertheless, emerging evidence has shown that several ATP synthase subunits are present on the plasma membrane (PM) of various cells, including endothelial cells, adipocytes, keratinocytes, hepatocytes, and some types of cancer cells (Arakaki *et al*, 2003; Inoue *et al*, 2007; Kim *et al*, 2004; Li *et al*, 2017; Yonally *et al*, 2006). Based on its ectopic location, this type of ATP synthase is referred to as ectopic ATP (eATP) synthase. Similar to the ATP synthase located on mitochondria, this cell-surface eATP synthase possesses catalytic activity that facilitates the generation of ATP in the extracellular environment to establish a suitable microenvironment. For instance, cancer cells under hypoxic conditions increase the catalytic activity of their eATP synthase to adapt to this unfavorable environment (Ma *et al*, 2010). In addition, ATP synthase subunit β (ATP5B), which is located on the PM, is a high-density lipoprotein receptor involved in the regulation of cholesterol homeostasis in HepG2 hepatocellular carcinoma cells (Martinez *et al*, 2003). eATP synthase is also expressed in antigen-presenting cells to facilitate the transportation of human immunodeficiency virus 1 (HIV-1) to immune cells (Yavlovich *et al*, 2012).

In the search for therapeutic interventions, an increasing number of studies have identified eATP synthase as a potential molecular target. The first finding supporting this possibility indicates that angiostatin, an antibody against ATP synthase subunits α and β, blocks the enzymatic activity of eATP synthase on the cell surface, with consequent suppression of proliferation, migration, and angiogenesis in endothelial cells (Moser *et al*, 2001). Since then, numerous PM ATP synthase inhibitors have been investigated (Hong *et al*, 2008). Treatment with these eATP synthase inhibitors reduces the production of extracellular ATP and leads to cytotoxic effects in neurons, adipocytes, and vascular endothelial cells (González-Pecchi *et al*, 2015; Xing *et al*, 2011). In addition to normal cells, eATP synthase blockade also inhibits the proliferation of various types of tumor cells (Fliedner *et al*, 2015; Lu *et al*, 2009; Mowery *et al*, 2008; Ravera *et al*, 2011). Similarly, our previous studies have revealed that the novel eATP synthase inhibitor citreoviridin induces cell cycle arrest and suppresses proliferation in both breast and lung cancer cells *in vitro* and *in vivo* (Chang *et al*, 2012; Chang *et al*, 2014; Hu *et al*, 2015; Wu *et al*, 2013).

Although eATP synthase has been studied for many years, its overall mechanism of transport to the cell surface remains unclear. In this study, we first performed spatial proteomic profiling to reveal the subcellular distribution of ATP synthase in cells. Then, transcriptome-based approaches were used to identify the mitochondrial dynamics and the microtubule-dependent motor protein involved in eATP synthase trafficking. Finally, a series of cellular methodologies, including super-resolution and real-time live-cell imaging, immunofluorescence, and flow cytometry were used to validate the eATP synthase trafficking pathway (Fig EV1).

## Results

### Real-time live imaging reveals that ATP synthase moves from the perinuclear region to the PM

Our previous studies have demonstrated the presence of eATP synthase on the cell surface of fixed cells using flow cytometry and immunofluorescence (Chang et al., 2012). However, there is limited real-time evidence regarding the dynamic spatial movement of ATP synthase. In this experiment, we attempted to monitor the subcellular trafficking of ATP synthase in live cells using real-time live imaging. Most fluorescent proteins are easily photobleached, making long-term observation difficult. Therefore, we manipulated photo-activatable green fluorescent protein (paGFP), which displays a 100-fold higher green fluorescence intensity and is more stable over several days than GFP (Karbowski *et al*, 2014). Thus, after photo-activation, the redistribution of activated paGFP could be followed over time. This permitted us to track the same targeted components that had been activated at the beginning of the experiment, instead of unlabeled, newly-synthesized components. First, we introduced the sequence of ATP5B, an enzymatic β subunit of the ATP synthase complex, into the paGFP vector (Fig EV2A). Next, we transfected this recombinant construction into lung cancer cells to observe the transport of ATP synthase. After stimulation with 405 nm light, the fusion of paGFP with ATP5B (ATP5B-paGFP) was recorded every 4 sec for 30 min (Fig EV2B). The microscopy images showed that the ATP5B-paGFP signal spread out after exposure to a pulse of 405 nm light (Fig EV2C). Consistently with this, western blotting with antibodies against GFP confirmed that cells transfected with the ATP5B-paGFP construct expressed the ATP5B-paGFP fusion protein (Fig EV2D). To identify the subcellular locations of ATP5B, CellMask Deep Red and MitoTracker Red were used to label the PM and mitochondria, respectively (Fig EV2E). Although fluorescence colocalization analysis indicated that approximately 98% of ATP5B-paGFP colocalized with the mitochondria (Fig EV2F-H), the time-series images showed that a substantial amount of ATP5B-paGFP was trafficked from the cytosolic region to the PM (Fig 1A and B). The process of ATP synthase trafficking from the cytosol to the PM took from a few seconds to several minutes.

**Figure 1.**
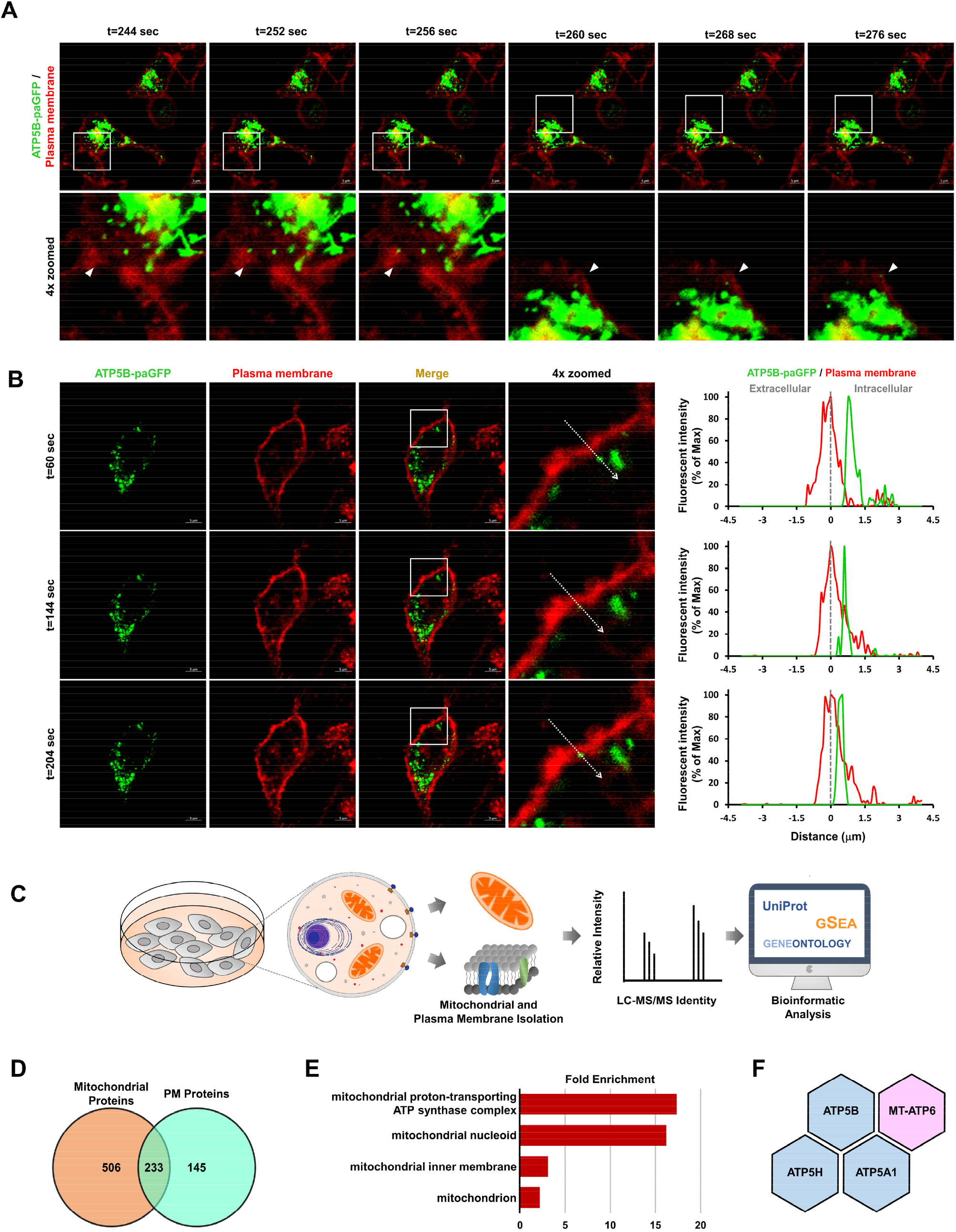
ATP synthase is routed to the cell surface via a mitochondria-dependent trafficking pathway. A Time-lapse images of ATP5B-paGFP signal (green) in live A549 cells were visualized using confocal microscopy (63× with 2× zoom). The plasma membrane was labeled using CellMask (1:10,000 dilution). Arrows show the colocalization of the ATP5B subunit and the PM with yellow fluorescence. The capture times after photoactivation with a 405 nm laser are shown. Scale bars, 5 μm. B Fluorescent signals of real-time tracing images are shown independently or merged. The fluorescence intensity profile for each channel (green and red represent ATP5B-paGFP and the PM, respectively) across the arrow on the enlarged image is shown in the line graph. Dotted lines indicate the plasma membrane. Scale bars, 5 μm. C Schematic representation of the cell surface and mitochondrial proteomes. D Venn diagram displaying the number of proteins in common between the mitochondria and PM of A549 cells. E Functional enrichment analysis of dual-localized proteins in mitochondria and PM. The terms were ranked according to fold enrichment, as shown on the *x*-axis. F The subunits of the ATP synthase complex that are common to both the mitochondria and the PM. The blue color represents the nucleus-encoded subunits, while the pink color represents the mitochondria-encoded subunit.

We then wanted to know whether the whole mitochondrial ATP synthase complex was routed to the PM. Therefore, analysis of the integrative organelle proteome was conducted to identify proteins that are commonly located on both mitochondria and the PM (Fig 1C). Altogether, 739 proteins were identified from purified mitochondria, and 378 were identified in the PM proteome (Fig 1D; Table EV1 and 2). Comparison of the mitochondrial and PM proteomes revealed 233 proteins that existed in both mitochondria and the PM (Fig 1D; Table EV3). The Gene Ontology (GO) enrichment analysis of these proteins showed that functions related to the mitochondrial proton-transporting ATP synthase complex were significantly enriched (Fig 1E), including ATP5B, ATP synthase subunit a (MT-ATP6), ATP synthase subunit d (ATP5H), and ATP synthase subunit α (ATP5A1) (Fig 1F). The existence of mitochondria-encoded subunit a, which lacks the sequence that targets the cell surface PM, implies that the ATP synthase complex may be assembled in the mitochondria and subsequently routed to the cell surface.

### Mitochondria act as delivery vehicles for the transport of assembled ATP synthase to the PM

In addition to the subunits of ATP synthase, other specific mitochondrial proteins (e.g., mitochondrial import receptor subunit TOM70 [TOMM70], malate dehydrogenase 2 [MDH2], and acetyl-CoA acetyltransferase [ACAT1]) were observed on the PM. Therefore, we speculated that the eATP synthase complex probably reaches the PM via mitochondria-dependent transportation. Based on this hypothesis, we conducted a bioinformatics analysis to validate the correlation between the expression of eATP synthase and mitochondrial trafficking-related genes. First, we measured the expression levels of eATP synthase via immunofluorescence microscopy and flow cytometry (Fig 2A-C). Next, we divided neuroblastoma cell lines into two groups based on their expression of eATP synthase. SK-N-BE(2)C and SK-N-DZ cells, which contained more eATP synthase, were defined as eATP synthase^high^ cells, whereas SK-N-AS and SK-N-SH cells were classified as eATP synthase^low^ cells (Fig 2D). The gene expression profiles of cells representing each group were curated from the Gene Expression Omnibus (GEO) with accession number GSE78061. Using a threshold of *p* < 0.05 and fold change ≥ 1.5, 496 significantly up-regulated and 605 significantly down-regulated differential genes were identified in eATP synthase^high^ versus eATP synthase^low^ cells (Fig EV3A; Table EV4). Subsequent functional enrichment analysis showed that many of the significantly differentially expressed genes are involved in mitochondrial organization and transportation (Fig EV3B; Table EV5). Consistent with this, gene set enrichment analysis (GSEA) revealed that the expression of genes associated with the set “mitochondrial transport” (GO: 0006839) was significantly elevated (*p* = 0.005) in eATP synthase^high^ than in eATP synthase^low^ cells (Fig 2E). In addition to neuroblastoma cell lines, we also found that genes associated with the following GO terms were significantly up-regulated in lung cancer cell lines in response to high expression of eATP synthase: “Mitochondrion transport along microtubule (GO: 0047497)”, “Establishment of mitochondrion localization, microtubule-mediated (GO: 0034643)”, and “Mitochondrion localization (GO: 0051646)” (Fig EV3C). The PM and mitochondria of cells transfected with ATP5B-paGFP were labeled using CellMask Deep Red and MitoTracker Red to observe the relationship between ATP synthase, mitochondria, and the PM. The time-series live-cell images showed that the ATP synthase β subunit was colocalized with mitochondria, and that they moved together toward the PM region (Fig 2F). We further speculated how ATP synthase in the mitochondrial inner membrane transfer to the PM by utilizing dual-color super-resolution microscopy, which provides a nanoscopic resolution to approach molecule dimensions of the proteins. The results showed that signals of TOM20 and the ATP synthase complex simultaneously presented on the mitochondria located near either the nucleus or PM, implying that whole mitochondria, containing the mitochondrial outer membrane (MOM) and the mitochondrial inner membrane (MIM), might be transported toward the cell membrane (Fig 2G). To further investigate the contact between the PM, MOM and MIM, we respectively labeled these three organelles by using the antibodies against their abundant proteins and observed their relative position via super-resolution imaging. TOM20, a component of mitochondrial import receptor complex located at the surface of the MOM, contacted with E-cadherin, a transmembrane glycoprotein of PM (Fig 2H). Also, the ATP synthase complex, an essential catalyst for ATP production located on the MIM, spatial colocalized with the Na^+^K^+^-pump, which is highly expressed on the PM (Fig 2I). These spatial colocalizations of mitochondrial membrane proteins to PM proteins might result from the fusion of the MOM and PM, and then the MIM and PM. Moreover, the colocalized ratio of TOM20 and E-cadherin has no difference with that of the ATP synthase complex and Na^+^/K^+^ pump (Fig 2J). Hence, we conclude that ATP synthase is assembled in mitochondria and then serves as the cargo of microtubule-dependent mitochondrial transport, and finally is transported to the cell surface by attachment of the MOM and MIM with PM in turn.

**Figure 2.**
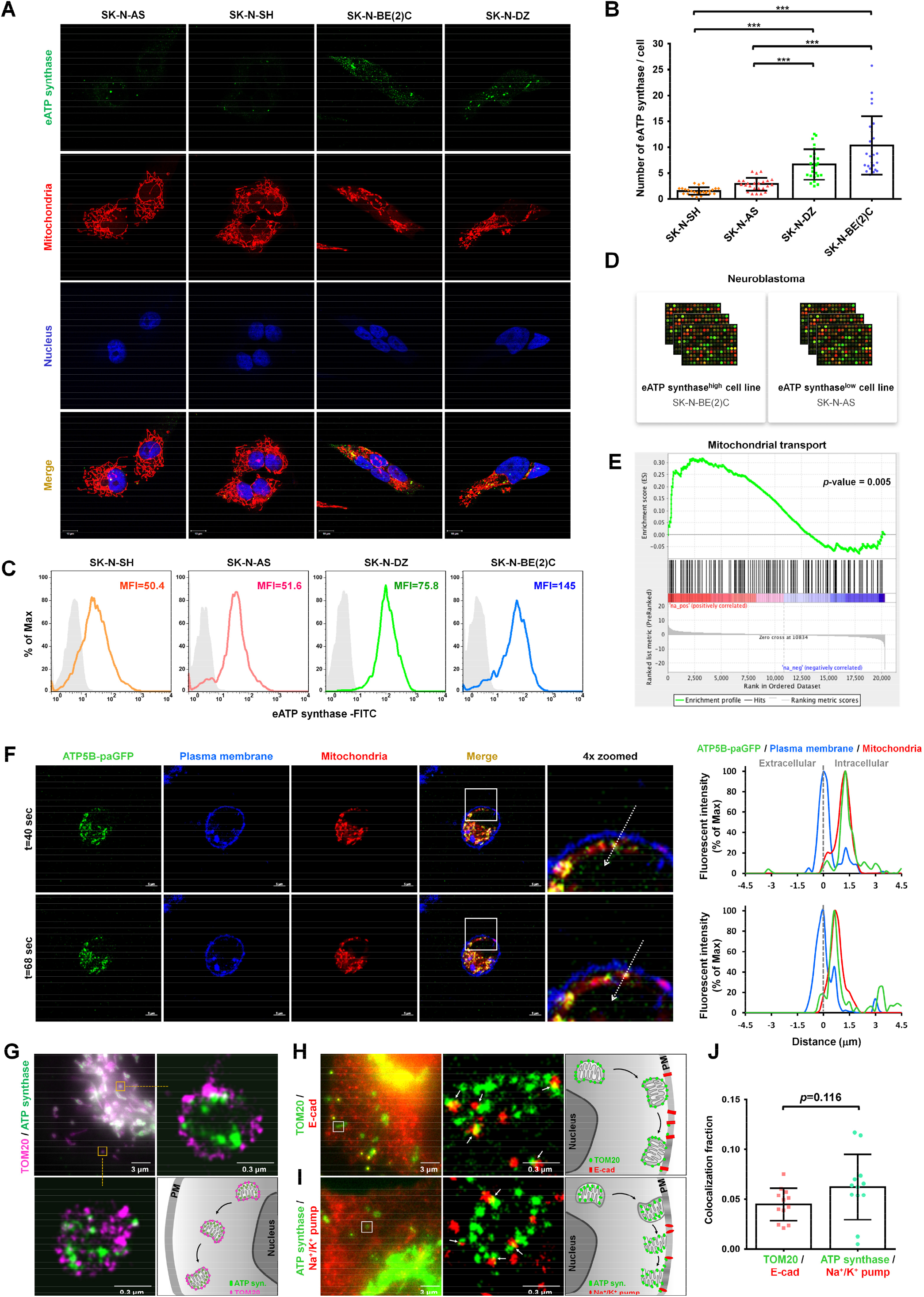
ATP synthase is ectopically translocated on the PM via mitochondria-dependent transport along the microtubules. A The abundance of eATP synthase on the cell surface of non-permeable neuroblastoma cells was detected using immunofluorescence. The antibody against the ATP synthase complex was used (green). Mitochondria were labeled using Mitotracker (1:10,000 dilution, red), nuclei were labeled using DAPI (blue). Fluorescence signals in the images are shown independently or merged. Scale bars, 10 μm. B Quantifications for eATP synthase counts per cell were measured using ImageJ software. Data presented are mean ± SD (n = 20). C The expression of eATP synthase on the cell surface was analyzed via flow cytometry, using antibodies for the ATP synthase complex or control mouse IgG. The mean fluorescence intensity (MFI) of eATP synthase based on the flow cytometry data is displayed. D The eATP synthase trafficking-related signature was categorized into two groups based on the expression of eATP synthase. E Gene set enrichment analysis (GSEA) was performed on the gene expression profiles of the eATP synthase^high^ and eATP synthase^low^ cell lines. The gene sets associated with mitochondrial transport were significantly positively enriched. F The fluorescence intensity profile for each channel (green, blue, and red represent ATP synthase β-paGFP, PM, and mitochondria, respectively) across the arrow on real-time tracing images is shown. Dotted lines indicate the plasma membrane. Scale bars, 5 μm. G dSTORM images show the distribution of TOM20 (magenta) and ATP synthase complex (green) at mitochondria in cells. H-I Two-color dSTROM images show the localization between these proteins. Cells were stained with the antibodies against TOM20 and E-cadherin (H) and ATP synthase complex and Na^+^-K^+^ pump (I). J The colocalization between the mitochondrial membrane and plasma membrane proteins were further quantified and shown as the bar chart. * *p* < 0.05, ** *p* < 0.01, *** *p* < 0.001.

### Mitochondrial dynamics influence the translocation of ATP synthase to the PM

Recent studies have found that mitochondrial dynamics such as fusion and fission is crucial to mitochondrial transport (Chen *et al*, 2009), and increasing the fission rate enhances the efficiency of mitochondrial transport (van der Bliek *et al*, 2013). Therefore, we investigated whether mitochondrial morphology affects the expression of eATP synthase, using the bio-image analysis software Icy (de Chaumont *et al*, 2012) (Fig 3A). Immunofluorescence revealed that mitochondria prefer fission in the eATP synthase^high^ cells A549 and SK-N-BE(2)C compared to the eATP synthase^low^ cells SK-N-AS (Fig 3B). Quantitative results showed that both the size and perimeter of mitochondria were negatively correlated with the abundance of eATP synthase (Fig 3C-E). Consistently, the protein and mRNA expression levels of GTPase dynamin-related protein 1 (DRP1), a crucial mediator of mitochondrial fragmentation (Palmer *et al*, 2013), were markedly higher in eATP synthase^high^ cells than in eATP synthase^low^ cells (Fig 3F and G). This observation suggests that the transport of eATP synthase may be dependent on the mitochondrial fission/fusion machinery.

**Figure 3.**
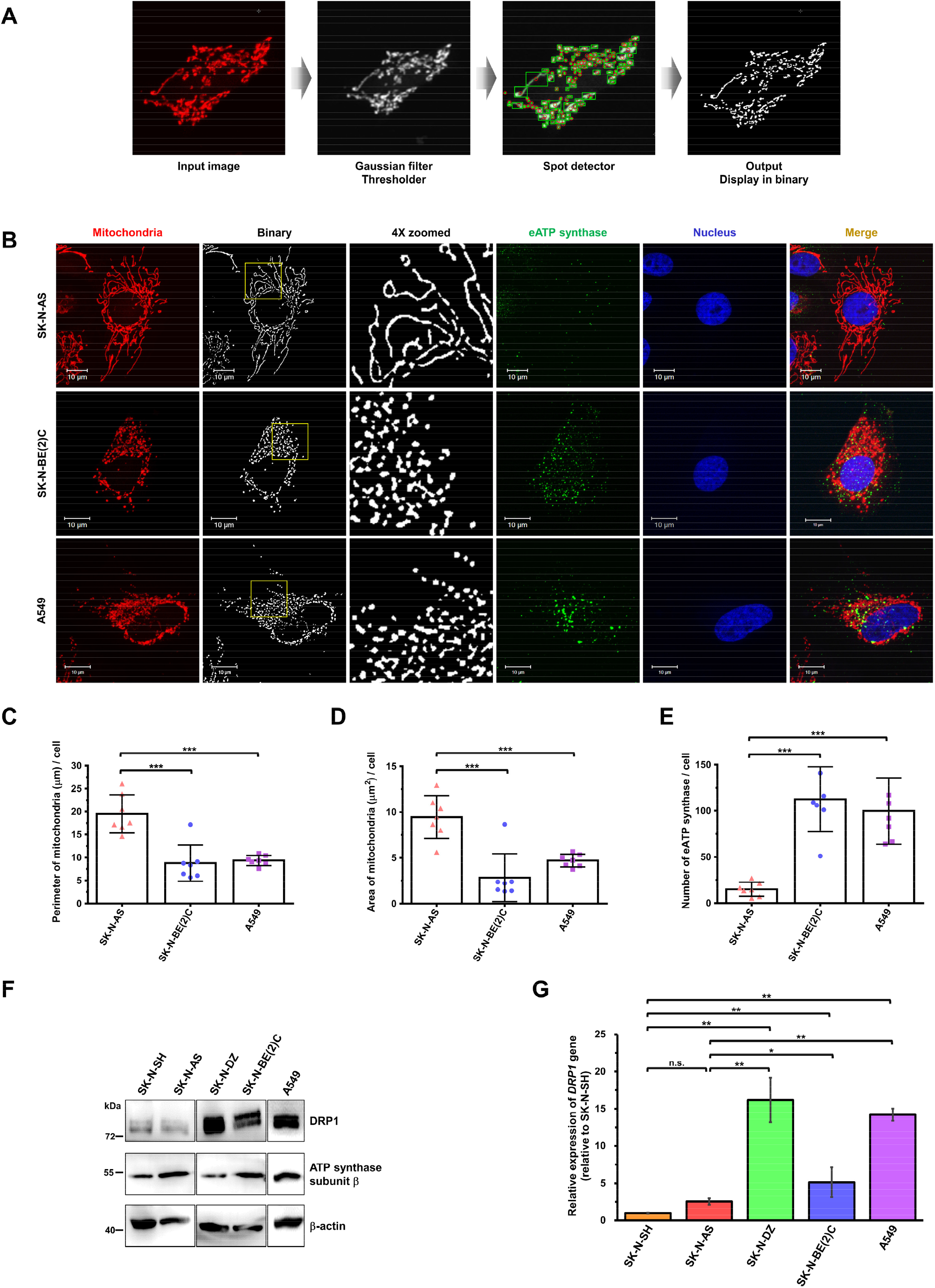
Mitochondrial dynamic morphology influences the expression level of ectopic ATP synthase. A Illustration of the process of quantifying the mitochondrial area using Icy software. B The mitochondrial morphology and the expression of eATP synthase in non-permeable neuroblastoma and lung cancer cells were visualized using anti-ATP synthase antibody followed by hybridization with Alexa 488 anti-mouse IgG (green) and Mitotracker (red) under confocal microscopy (100×). Nuclei were labeled using DAPI (blue). Binary images were processed using Icy software (lower). Scale bars, 10 μm. C–E Quantification of the mitochondrial perimeter (C) and area (D) by Icy software, and of the eATP synthase count (E) by ImageJ software (n = 7). Data presented are the mean ± SD. F The expression levels of ATP synthase subunit β and DRP1 in various cell lines were detected using western blotting. G The gene expression levels of DRP1 were measured via quantitative RT-PCR in various cell lines. Values were normalized to the expression of GAPDH and relative to the expression of the DRP1 gene in SK-N-SH cells. Values shown are the mean ± SEM (n = 3) * *p* < 0.05, ** *p* < 0.01, *** *p* < 0.001.

### DRP1-dependent mitochondrial fission regulates the anterograde transport of ATP synthase

We wished to gain more insight into the relationship between mitochondrial dynamics and eATP synthase transport. Mitochondrial division inhibitor 1 (Mdivi-1) was used to explore whether attenuation of mitochondrial fission affects the expression of eATP synthase. As expected, treatment with Mdivi-1 for 24 h resulted in the elongation of mitochondria (Fig EV4A). We then examined whether this affected the abundance of eATP synthase on the cell surface of eATP synthase^high^ cells. Visualized immunofluorescence and flow cytometry indicated that the expression of eATP synthase was down-regulated after treatment with Mdivi-1 (Fig EV4A-C). The protein levels of Drp1 were significantly reduced after treatment with Mdivi-1 in the eATP synthase^high^ cell lines A549 and SK-N-BE(2)C (Fig EV4D). The real-time movement of ATP5B-pa-GFP was recorded to confirm this finding. Time-series images demonstrated that ATP5B-pa-GFP moved a shorter distance in Mdivi-1-treated cells than in control cells (Fig EV4E and F). Fluorescence colocalization analysis showed that the overlapping proportion of ATP5B-pa-GFP and Deep Red-labeled PM was reduced in Mdivi-1-treated cells (Fig EV4G and H).

These results indicate that disruption of mitochondrial dynamics modulates the transport of ATP synthase from the cytosol to the PM. In addition, silencing of DRP1 to disrupt mitochondrial dynamics was performed in eATP synthase^high^ cells to address the specific regulation of ATP synthase transport by mitochondrial fission. The expression of DRP1 protein in A549 and SK-N-BE(2)C cells transfected with short interfering RNA (siRNA) for DRP1 was severely reduced (Fig 4A). These DRP1-depleted cells exhibited decreased eATP synthase on the cell surface (Fig 4B-D). In comparison, in the eATP synthase^low^ cells SK-N-AS, overexpression of DRP1 (Fig 4E) activated mitochondrial division and elevated the expression of eATP synthase on the cell surface (Fig 4F-H). Combined, these results indicate that DRP1 is required for the transport of ATP synthase from the cytosol to the PM.

**Figure 4.**
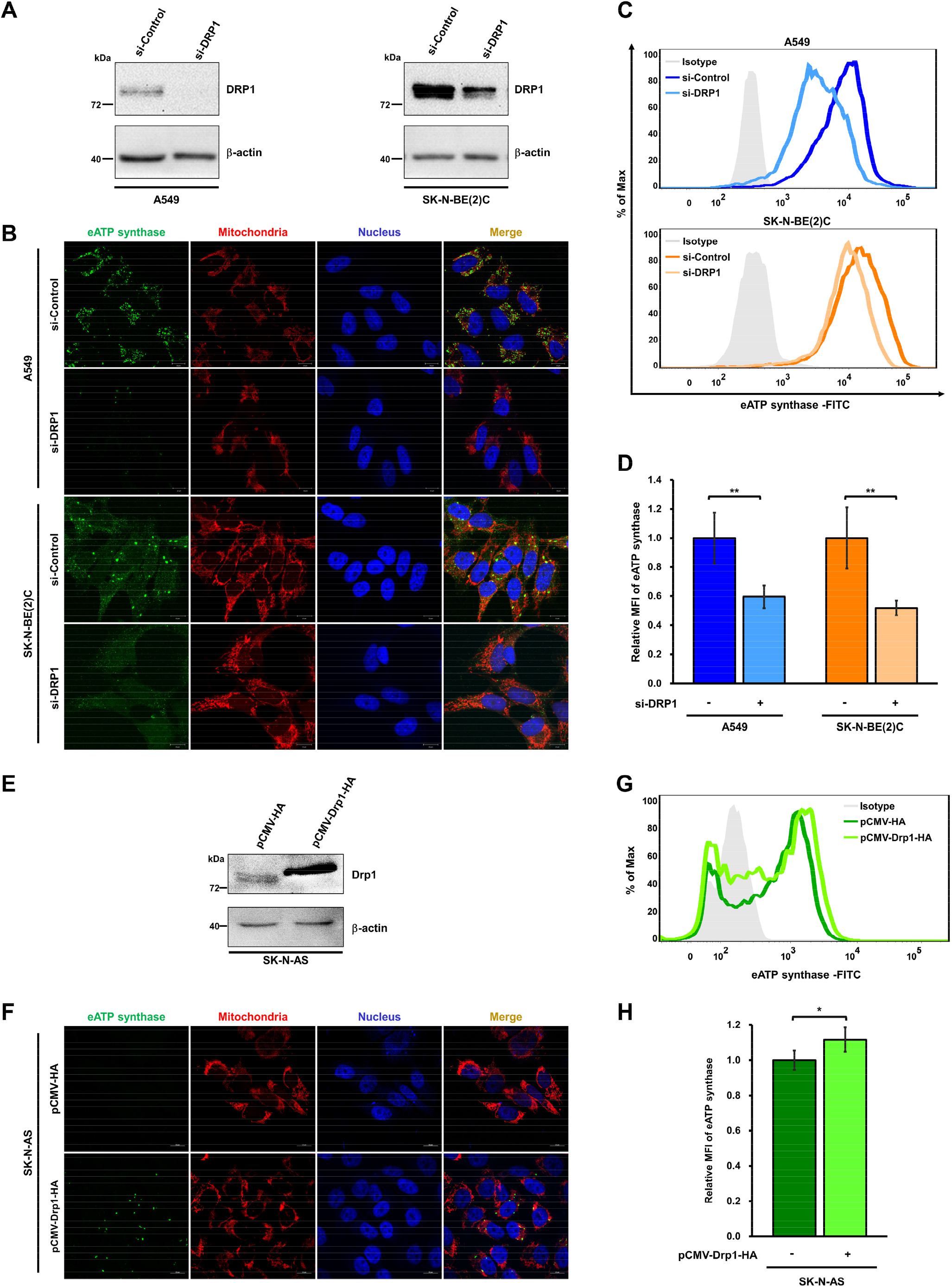
Disturbance of DRP1-dependent mitochondrial dynamics affects the trafficking of ectopic ATP synthase. A The knockdown efficiency of siDRP1 was measured via western blotting using anti-DRP1 antibody in siControl and siDRP1-transfected cells. B-D The expression of eATP synthase on the cell surface was detected via immunofluorescence (B) and flow cytometry (C) using anti-ATP synthase antibody in the siControl and siDRP1 groups without permeabilization. The mean fluorescence intensity (MFI) of eATP synthase from (C) is displayed as a bar chart (D). E The overexpression efficiency was measured via western blotting using anti-HA antibody in the control and HA-DRP1-overexpressing cells. F-H The expression of eATP synthase on the cell surface was detected via immunofluorescence (F) and flow cytometry (G) using anti-ATP synthase antibody in the HA-control and HA-DRP1-overexpressing groups without permeabilization. The relative MFI of eATP synthase from (F) is displayed as a bar chart (H). Values shown are the mean ± SD (n = 3). * *p* < 0.05, ** *p* < 0.01, *** *p* < 0.001.

### Microtubules act as the intracellular railroad for ectopic ATP synthase trafficking

Previous research has shown that in mammalian cells, mitochondrial transport occurs along microtubule tracks (López-Doménech *et al*, 2018), which is consistent with the GSEA results of the present study. Therefore, we hypothesized that microtubules play a critical role in mitochondria-dependent eATP synthase trafficking. Before testing this hypothesis, we used super-resolution imaging to dissect whether mitochondria localize near microtubules and the results illustrated that the puncta of TOM20, located at the outer membrane of mitochondria, was observed near microtubules (Fig 5A). To further understand whether microtubules act as the intracellular railroad for ectopic ATP synthase trafficking, nocodazole (a microtubule depolymerization agent) was used to disrupt the microtubules in eATP synthase^high^ cells. After treatment with nocodazole for 1 h, the microtubules were depolymerized and the abundance of eATP synthase was reduced (Fig 5B). Flow cytometry also showed that nocodazole inhibited the expression of eATP synthase on the cell surface of both A549 and SK-N-BE(2)C cells (Fig 5C and D). In addition, time-lapse images revealed that the movement of ATP5B-pa-GFP was reduced in nocodazole-treated cells versus in control cells (Fig 5E and F). The colocalization analysis showed that ectopic ATP5B-pa-GFP, which colocalized with the PM, was also reduced following treatment with nocodazole (Fig 5G and H). These results illustrate that microtubules are involved in mitochondria-dependent ATP synthase trafficking from the cytosol to the cell surface.

**Figure 5.**
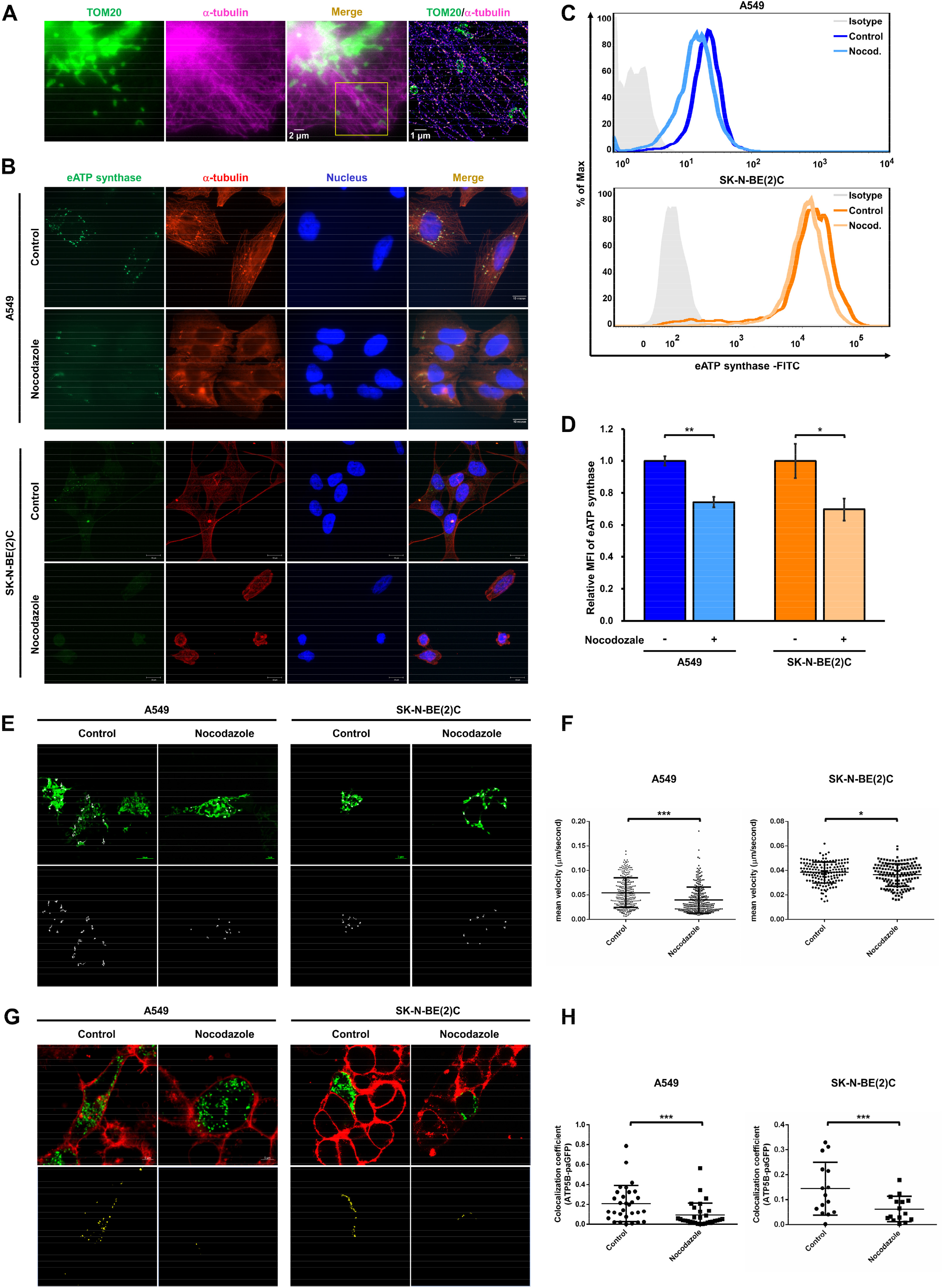
Microtubule-dependent mitochondrial transport regulates the transport of ectopic ATP synthase. A Cells were stained with TOM20 (green) and α-tubulin (magenta) to confirm their localization. 100×oil-immersion objective was used for wide-field illumination. Yellow box shows the super-resolution image captured by dSTORM system. B Both A549 and SK-N-BE(2)C cells were treated with either DMSO or nocodazole (25 μM) to disrupt their microtubules. The expression of eATP synthase on the cell surface was detected using anti-ATP synthase antibody followed by hybridization with Alexa 488 anti-mouse IgG (green) in non-permeable cells. After probing eATP synthase on the cell surface, the cells were subjected into permeabilization and then the microtubules were labeling using anti-α-tubulin antibody, followed by hybridization with Alexa 555 anti-mouse IgG (red). Scale bars, 10 μm. C The expression of eATP synthase on the cell surface was detected via flow cytometry using anti-ATP synthase antibody in the DMSO- or nocodazole-treated cells. D The bar chart shows the relative MFI of eATP synthase. Data presented are the mean ± SD (n = 3). E The movement of the ATP synthase β subunit-paGFP fusion protein in the DMSO and nocodazole groups was recorded for 15 min via confocal microscopy (upper). Their tracks are shown as white lines and were processed using Metamorph software (lower). F The dot plot represents the quantification of the translocation velocity in the DMSO and nocodazole groups (n = 30). G The localization of the ATP synthase β subunit-paGFP fusion protein (green) and the PM (CellMask; red) was determined using confocal microscopy. The colocalization of these two signals is shown as yellow fluorescence in the merged images (upper), and was further processed using ZEN software (lower). H The dot plot shows the colocalization coefficient (colocalization fluorescent area versus total ATP synthase β subunit-paGFP fluorescent area) (n = 30). Values shown are the mean ± SD. * *p* < 0.05, ** *p* < 0.01, *** *p* < 0.001.

### Kinesin family member 5B (KIF5B) drives the anterograde transport of ATP synthase via the formation of a complex with DRP1

Kinesin family member 5B (KIF5B, also known as kinesin-1 heavy chain, KINH) is a driver of mitochondrial transport along microtubules (Reis *et al*, 2009). Hence, we examined the relationship between KIF5B and eATP synthase using the siRNA-mediated KIF5B gene knockdown approach. A comparison of siKIF5B and siControl-transfected cells showed that the level of KIF5B protein was lower in the former group (Fig 6A). Both immunofluorescence microscopy and flow cytometry showed that the abundance of eATP synthase was significantly down-regulated in KIF5B-depleted cells (Fig 6B-D). Based on these results, we inferred that KIF5B was responsible for the transport of eATP synthase.

**Figure 6.**
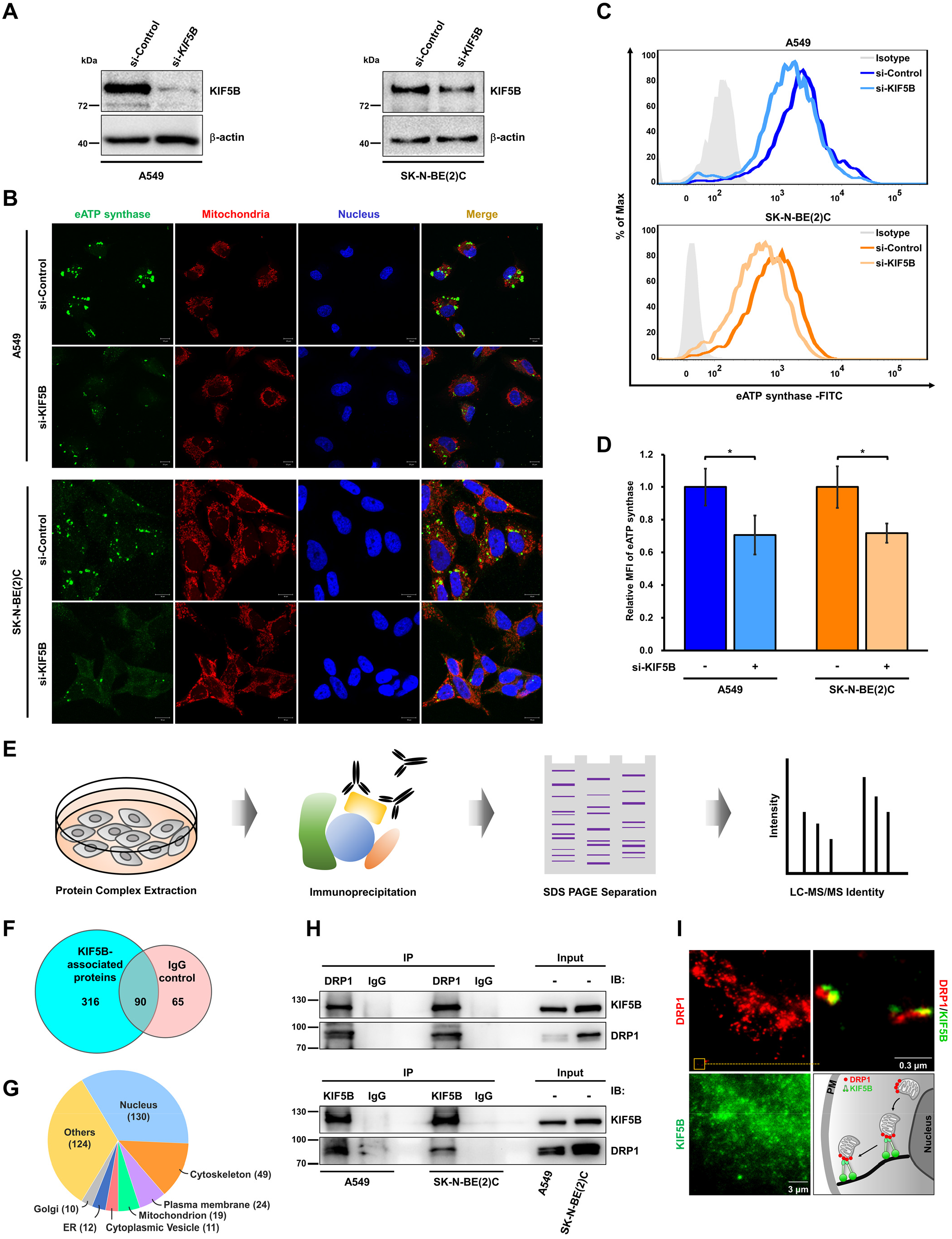
KIF5B and DRP1 form a transport complex to mediate intercellular trafficking of ATP synthase. A A549 and SK-N-BE(2)C cells were transfected with siControl or siKIF5B. The knockdown efficiency of siRNA was measured via western blotting using anti-kinesin-1 heavy-chain antibody. B-D The expression of eATP synthase on the cell surface was detected via immunofluorescence (B) and flow cytometry (C) using anti-ATP synthase antibody in siControl and siKIF5B-knockdown cells. The relative MFI of eATP synthase from C is displayed as a bar chart (D). Data presented are the mean ± SD (n = 3). * *p* < 0.05, ** *p* < 0.01. *** *p* < 0.001. E Schematic representation of interactomic profiling using anti-KIF5B or anti-DRP1 antibody. F The number of proteins pulled down by the KIF5B-interactome are shown as a Venn diagram. Anti-IgG antibody was used as the internal control of immunoprecipitation. G The subcellular localization of KIF5B-interacting proteins. The numbers represent the number of the proteins in the indicated subcellular locations. H Immunoprecipitation with anti-KIF5B antibody or anti-DRP1, followed by western blotting using the indicated antibodies, was performed to verify the interaction between KIF5B and DRP1. Representative images obtained from three independent experiments are shown. I Cells were stained with DRP1 (red) and KIF5B (green). The localization of DRP1 and KIF5B were further detected by two-color dSTORM imaging system. The diagram shows the relationship between mitochondria, DRP1 and KIF5B.

Nevertheless, the mechanism through which KIF5B interacts with mitochondria remains unclear. KIF5B-interactomic profiling (i.e., immunoprecipitation of KIF5B followed by mass spectrometry analysis) was used to systematically identify KIF5B-interacting proteins to investigate this mechanism (Fig 6E). A total of 316 proteins were identified in A549 cells (Fig 6F; Table EV6) and further classified according to their cellular locations (Fig 6G; Table EV7). Nineteen of these proteins were located in mitochondria. Interestingly, DRP1, which affects eATP synthase trafficking via regulation of mitochondrial fission, was identified as one of the KIF5B-interacting proteins. We further validated the interaction between KIF5B and DRP1 in A549 and SK-N-BE(2)C cells by co-immunoprecipitation, using antibodies against KIF5B and DRP1 (Fig 6H). Dual-color super-resolution imaging further revealed the spatial colocalization of DRP1and KIF5B (Fig 6I). As we know, DRP1 is generally localized on the mitochondria, and the results show that DRP1 also localizes around the cell surface with KIF5B. In conclusion, KIF5B plays an essential role in the trafficking of eATP synthase via the formation of a complex with DRP1.

### The variable domain of DRP1 plays a critical role for ectopic ATP synthase trafficking via interacting with KIF5B

To further understand how DRP1 interacts with KIF5B, the ClusPro 2.0 web server (https://cluspro.org) was used for protein-protein docking simulation (Kozakov *et al*, 2017). As shown in Fig 7A and B, structure-based domain arrangements presented the localization of conserved domains of DRP1 (Protein Data Bank code 4BEJ), including four major domains: the N-terminal GTPase domain, middle assembly domain, variable domain (VD), and C-terminal GTPase effector domain (GED) (Adachi *et al*, 2016). Since the crystal structure of the whole KIF5B protein is not available yet, we carried out homology modeling on the full-length sequence of KIF5B obtained from the UniProt database to build a protein structure using SWISS-MODEL (Waterhouse *et al*, 2018). Mitochondrial fission protein 1 (FIS1), which has been reported to recruit DRP1 to mitochondria and mediate mitochondrial fission by forming the FIS1-DRP1 complex (Chen *et al*, 2005), was used as a positive control of the protein-protein docking simulation. The interfaces for KIF5B and for FIS1 were located in two different regions of DRP1 (Fig 7C). The area for KIF5B-DRP1 interaction containing 10 residues (purple; ASN-511, GLU514, GLN-515, ARG-516, ASN-518, ARG-519, ARG-522, GLU-523, SER-526, ARG-530) is located on the VD of DRP1 (Fig 7D; Zoom2). Interestingly, six residues (violet; LYS-133, ASN-141, ASP-161, GLU-168, HIS-295, ASP-299), responsible for the interaction of FIS1 and DRP1, are located on the N-terminal GTPase domain of DRP1 (Fig 7D; Zoom1). To verify the results of the protein-protein docking simulation, full-length and C-terminal truncated DRP1 constructs (containing VD and GED), which were tagged with HA-tag, were generated (Fig 7E). Co-immunoprecipitation using antibodies against KIF5B indicated that the C-terminal of DRP1 was sufficient for the interaction of KIF5B with DRP1 (Fig 7F). Both immunofluorescence microscopy and flow cytometry showed that the abundance of eATP synthase was significantly up-regulated in SK-N-AS cells overexpressing full-length and C-terminal truncated DRP1 (Fig 7G-I). These results suggest that DRP1 regulates the trafficking of ectopic ATP synthase via the interaction of its variable domain with KIF5B (Fig 7J).

**Figure 7.**
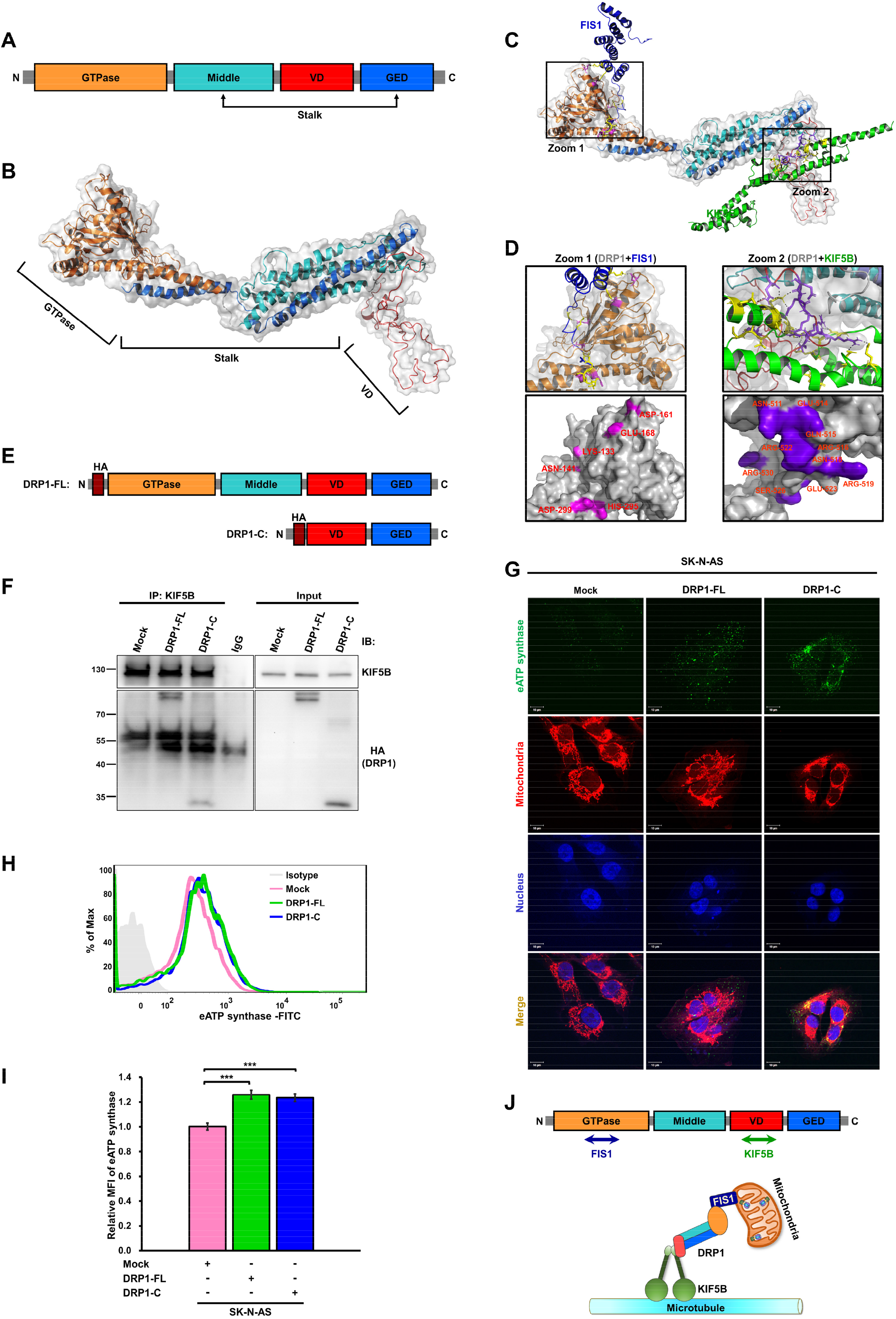
The variable domain of DRP1 is important for the interaction of DRP1-KIF5B complex regulating the transport of ATP synthase toward the cell surface. A Domain organization of DRP1 including N-terminal GTPase domain (orange), variable domain (VD, red), and stalk domain which is consist of middle domain (cyan) and GTPase effector domain (GED, blue). B Ribbon and transparent-surface representations show the tertiary structure of DRP1 (PDB ID: 4BEJ) and highlight the conserved domain indicated in top panel. C Overview of FIS1-DRP1 (navy) and KIF5B-DRP1 (green) interfaces. D Close-up views of FIS1-DRP1 (Zoom 1) and KIF5B-DRP1 (Zoom 2) interfaces show the two interaction sites, respectively. The residues involved in the interactions are represented as sticks, and hydrogen-bonding interactions are depicted as dashed lines (upper). Surface representation of two interacting sites in DRP1, with purple color indicating KIF5B-interacting sites and violet indicating FIS1-interacting sites. Interface residues are labelled (lower). E Scheme of the DRP1-HA truncated constructs used for immunoprecipitation. F Immunoprecipitation with anti-KIF5B antibody, followed by western blotting using anti-HA antibody, was performed to confirm the interaction between KIF5B and the c-terminal region of DRP1. G-I The expression of eATP synthase on the cell surface was detected via immunofluorescence (G) and flow cytometry (H) using anti-ATP synthase antibody in SK-N-AS cells transfected with the indicated DRP1 constructions. The relative MFI of eATP synthase based on the flow cytometry data is displayed as a bar chart (I). Data presented are the mean ± SD (n = 3). J Model for KIF5B-DRP1 complex which is responsible for transport of mitochondria. Firstly, DRP1 is recruited to the mitochondria by FIS1 known as a mitochondrial outer membrane protein and interacts to FIS1 with N-terminal GTPase domain (orange). After mitochondria fission regulating via FIS1-DRP1 complex, DRP1 leads the fragmented mitochondria to the microtubules by interacting to the motor protein, KIF5B, with variable domain (red). KIF5B-DRP1 complex subsequently acts as a vehicle for mitochondria transportation. * *p* < 0.05, ** *p* < 0.01. *** *p* < 0.001.

## Discussion

Numerous studies have revealed the roles of PM-located ATP synthase (eATP synthase) in cancers (Li et al., 2017; Lu et al., 2009; Speransky *et al*, 2019). In this work, we aimed to establish the mechanism responsible for the transport of eATP synthase. In our eATP synthase transport model, ATP synthase subunits, including mitochondria- and nucleus-encoded subunits, are already assembled in the mitochondria. Later, DRP1 promotes mitochondrial fission, which makes mitochondrial movement possible. Moreover, in association with KIF5B, DRP1 carries these fragmented mitochondria containing ATP synthase along the microtubules toward the cell surface. Some evidence points to a positive correlation between eATP synthase and the migratory capability of tumors (Li et al., 2017; Speransky et al., 2019). Identification of these ATP synthase trafficking-related regulators may thus provide useful information for the development of potential therapies against metastatic cancer.

Few studies have explored the question of how ATP synthase reaches the cell surface. To function properly, proteins in eukaryotic cells are usually transported to the appropriate destinations as encoded by short sequences of amino acid peptides, termed signal peptides or targeting sequences. The post-translational translocation pathway transports these proteins directly to the appropriate locations, such as the nucleus, mitochondria, or cytoplasm, via targeting sequences (Cokol *et al*, 2000; Neupert, 1997). For instance, proteins with mitochondria-targeting sequences (MTS) are delivered by cytosolic chaperones to the translocase of the outer membrane (TOM) receptors and imported into the mitochondrion by the translocase of the inner membrane (TIM) (Chacinska *et al*, 2009; Gabriel *et al*, 2003). There are no studies providing evidence regarding the presence of PM-targeting sequences in ATP synthase subunits. It is improbable that these subunits are recruited directly to the PM (Ma *et al*., 2010). In addition, Rai et al. detected eATP synthase on the cell surface of hepatocytes that contained both nucleus- and mitochondria-encoded subunits (Rai *et al*, 2013). Consistent with this, our work verified the presence of mitochondria-encoded subunit a, which lacks the PM-targeting sequence, on the cell surface (Fig 1F). Moreover, our bioinformatics analysis and real-time live imaging revealed that ATP synthase is transported from the mitochondria to the PM (Fig 2D-F and EV2). This evidence supports our hypothesis that the entire ATP synthase complex is assembled in the mitochondria and then delivered to the cell surface. However, this raises the seminal question of whether there is a possibility of fusion between the mitochondrial membrane and the PM. We used a super-resolution microscope to further reveal whether the mitochondrial membrane attaches to the PM. The results showed that the mitochondrial outer and inner membrane are close to the plasma membrane in turn to anchor ATP syntheses on the cell surface (Fig 2G-J). Recent studies have revealed that in budding yeast, a nuclear migration protein (Num1) tethers mitochondria to inner side of the PM and promotes fusion between these two cellular architectures (Ping *et al*, 2016). Thus, we suggest that in mammalian cells, mitochondria and the PM may fuse via the similar machinery, but this hypothesis requires further investigation.

Mitochondrial morphology is constantly changing between the fission and fusion states, and is influenced by metabolic and pathogenic conditions, as these affect the inner and outer environments of the mitochondria (Wikstrom *et al*, 2014). This dynamic state allows mitochondria to communicate with each other to carry out their ordinary functions (Chan, 2006). Disorders of mitochondrial dynamics lead to a range of disease pathologies (Amiott *et al*, 2008; Archer, 2013). Fusion produces hyper-tubulated forms to enhance communication between interconnected mitochondria. This interconnected mitochondrial network allows mitochondria to complement each other’s deficiencies to promote the health of mitochondria. In contrast, fission creates more fragmented mitochondria, which enhances the oxygen-generation rate and accelerates mitochondrial transport (Youle *et al*, 2012). Accumulating evidence suggests that in cancers, mitochondrial dynamics are imbalanced, with increased fission and decreased fusion (Caino *et al*, 2016; Vyas *et al*, 2016). Mitochondrial fission is mediated by DRP1, which oligomerizes to a ring-like structure to facilitate the scission of mitochondria (Mishra *et al*, 2014; Wai *et al*, 2016). Recent studies have determined that DRP1-dependent mitochondrial fission is crucial for the specific subcellular localization of mitochondria (Adachi et al., 2016; Favaro *et al*, 2019; Ishihara *et al*, 2009). Therefore, based on our transcriptomics analysis and these previous reports, we can infer the relationship between mitochondrial fragmentation and the abundance of ATP synthase. Indeed, our experiments show that a fragmented mitochondrial network enhances the expression of eATP synthase in neuroblastoma cell lines (Fig 3B-E). Additionally, we disrupted mitochondrial fission via interfering with DRP1 expression (Fig 4 and EV3). The results showed that this disruption triggered mitochondrial fission and affected the abundance of eATP synthase on the cell surface.

As with most cargos, mitochondrial transport is accomplished by motor proteins, always including kinesin, working along the microtubules by changing their conformation involving ATP hydrolysis (Cai *et al*, 2016). Treatment with microtubule-targeting drugs significantly inhibits the microtubule-related organelle transport velocity of mitochondria in human neuroblastoma cells (Smith *et al*, 2016). The results of our study demonstrate that disruption of microtubule polymerization suppresses the expression level and translocation velocity of eATP synthase (Fig 5). Generally, kinesin physically attaches itself to the transported cargo via the adaptor protein, which acts as a linker between the motor protein and the cargo-surface protein (Barlan *et al*, 2017; Isojima *et al*, 2016). In neural cells, mitochondria are transported along microtubules by a motor complex composed of KIIF5B, trafficking kinesin-binding protein 1 (TRAK1) and mitochondrial Rho GTPase (Miro) (Mishra & Chan, 2014). Tanaka et al. found that, in *kif5B^−/−^* mice, mitochondria were clustered at the minus-ends of microtubules and accumulated around the nucleus in neural cells (Tanaka *et al*, 1998). Therefore, we speculate that KIF5B may play a critical role in eATP synthase trafficking by mediating mitochondria-dependent transport. Indeed, our immunofluorescence and flow cytometry analyses demonstrated that the abundance of eATP synthase was reduced as a result of silencing KIF5B (Fig 6A-D). By combining co-immunoprecipitation and liquid chromatography–tandem mass spectrometry (LC–MS/MS) spectra, we also noted that the mitochondrial outer membrane protein DRP1 associated with KIF5B to transport ATP synthase toward the cell surface (Fig 6E-I).

In conclusion, we have identified a novel transport mechanism for eATP synthase, in which DRP1 leads to mitochondrial fission and subsequently associates with KIF5B to deliver the assembled ATP synthase complex within fragmented mitochondria along microtubules to the cell surface. The mitochondrial outer membrane and inner membrane attach to the plasma membrane in turn to anchor ATP syntheses on the cell surface.

## Materials and Methods

### Cell cultures

Human lung cancer cell line A549 and neuroblastoma cell lines SK-N-DZ, SK-N-BE(2)C, SK-N-SH, and SK-N-AS were purchased from the American Type Tissue Collection (Manassas, VA, USA). All cell lines were cultured in Dulbecco’s modified Eagle medium (DMEM; Thermo Fisher Scientific, Waltham, MA, USA) supplemented with 10% fetal bovine serum (Gibco, Thermo Fisher Scientific, Waltham, MA, USA), in a 37°C incubator with 5% carbon dioxide. Cells were routinely passaged when they reached 80–90% confluency. All cells were detached using trypsin/ethylenediaminetetraacetic acid (trypsin/EDTA) solution (Thermo Fisher Scientific) and transferred to new culture dish for further incubation.

### Reagents and antibodies

All chemicals used in this study were dissolved in dimethyl sulfoxide (DMSO; Sigma–Aldrich, St. Louis, MO, USA). Mdivi-1 (Sigma–Aldrich) was solubilized at 50 mM and nocodazole (Sigma–Aldrich) was solubilized at 33 mM as stocks. The stocks were stored at −20°C and freshly diluted in culture medium at the indicated concentration for treatment. The control samples were treated with an equal volume of DMSO only. The mitochondria in cells were stained with MitotrackerRed (Thermo Fisher Scientific) at a 1:10,000 dilution in serum-free medium at 37°C for 20 min. The PM was stained with CellMask (Thermo Fisher Scientific) at a 1:10,000 dilution in phosphate-buffered saline (PBS) at 37°C for 10 min. The primary antibodies against the following proteins were used in this study: ATP synthase complex (Abcam, Cambridge, MA, USA), KIF5B (Abcam) and DRP1 (Abcam), α-tubulin (GeneTex, Irvine, CA, USA), ATP synthase β subunit (GeneTex), HA-tag (BioLegend, San Diego, CA, USA), and β-actin (Sigma–Aldrich).

### Organelle proteomic profiling

A549 lung cancer cells (2 × 10^7^) were washed twice with cold PBS. PM proteins were extracted using a cell surface isolation kit (Thermo Fisher Scientific), whereas mitochondrial proteins were isolated using the Mitochondria Isolation Kit (Thermo Fisher Scientific) according to the instructions provided by the manufacturer. Total proteins from diverse organelles were extracted using a phase-transfer surfactant solution containing 12 mM sodium deoxycholate (Sigma–Aldrich), 12 mM sodium lauroyl sarcosine (MP Biomedical, Santa Ana, CA, USA), and 100 mM triethylammonium bicarbonate (TEAB; Sigma–Aldrich). A protease inhibitor cocktail (BioShop, Burlington, Canada) was used to prevent protein degradation. The mixtures were sonicated on ice with a 0.6 s cycle at 60% amplitude and centrifuged at 16,000 × g for 20 min at 4°C. The concentration of proteins was measured using an enzyme-linked immunosorbent assay reader (BioRad, Hercules, CA, USA) at 570 nm using the Pierce BCA Protein Assay Kit (Thermo Fisher Scientific). A total of 50 μg proteins from each organelle underwent reduction with dithiothreitol/50 mM TEAB at room temperature for 30 min, and alkylation with iodoacetamide/50 mM TEAB at room temperature for 30 min in the dark. Alkylated proteins were digested by Lys-C (1:100 w/w) (WAKO, Richmond, VA, USA) at 37°C for 3 h and trypsin (1:100 w/w) (Thermo Fisher Scientific) at 37°C for 16 h. The digested peptides were acidified with trifluoroacetic acid (TFA, Sigma–Aldrich) until the pH was < 3, and then treated with ethyl acetate (Sigma–Aldrich) in 10% TFA to remove detergents. After vacuum-drying using a SpeedVac (EYELA, Tokyo, Japan), the peptides were dissolved with 0.1% (v/v) TFA and 5% (v/v) acetonitrile (Thermo Fisher Scientific) and subjected to desalting using StageTips with SDB-XC Empore disk membranes (SDB-XC StageTip; 3M, St. Paul, MN, USA)(Rappsilber *et al*, 2007).

### Immunoprecipitation

After washing twice with PBS, the cells were lysed using Pierce IP lysis buffer (Thermo Fisher Scientific) with protease inhibitor (BioShop) to repress protein degradation. A total of 3 mg of Dynabeads Protein A (Thermo Fisher Scientific) were blocked with 5% bovine serum albumin (BioShop, Ontario, Canada) at 4°C for 30 min and incubated with 1 mg cell lysate at 4°C for 1 h. After precleaning, the precleared cell lysate was further hybridized with 2 μg primary antibody at 4°C overnight. Anti-IgG antibody (Abcam) was utilized as the negative control. Protein– antibody complexes were incubated with Dynabeads Protein A at 4°C for 2 h. The targeted protein-interacting partners were further analyzed by western blotting and nano-LC–MS/MS analysis.

### Nano-LC–MS/MS analysis and proteomic data analysis

The peptides were identified via nano-LC–MS/MS using an LTQ-Orbitrap XL (Thermo Electron, Bremen, Germany) equipped with a nanoACQUITY ultra-performance liquid chromatography system (Waters, Milford, MA, USA) as previously described (Hu et al., 2015). The raw MS spectra data were analyzed using MaxQuant software (Cox *et al*, 2008) for peak detection and protein identification (version 1.3.0.5. for PM proteins; version 1.5.2.8 for mitochondrial proteins and co-immunoprecipitation), using the Swiss-Prot annotated and reviewed database (Cox *et al*, 2011) (published in 2014 for PM proteins, and in 2016 for mitochondrial proteins and co-immunoprecipitation). The parameter settings for the search criteria were identical to those employed in our previous study (Hu et al., 2015). The identified proteins were analyzed using the Database for Annotation, Visualization, and Integrated Discovery (http://david.abcc.ncifcrf.gov/tools.jsp) with default settings, and we focused only on the biological process category.

### Gene expression analysis

The gene expression profiles used in this study were mined from GEO (https://www.ncbi.nlm.nih.gov/geo/) using the R package “GEOquery” with accession number GSE78061 for the neuroblastoma cell lines SK-N-DZ, SK-N-BE(2)C, SK-N-AS, and SK-N-SH. According to their expression levels of ectopic ATP synthase determined through flow cytometry, the cell lines were separated into two groups, eATP synthase^high^ and eATP synthase^low^ cells. To avoid the platform effect, we considered only the gene expression profiles provided in Affymetrix UGU 133 plus 2.0, and the raw expression datasets were normalized using the frozen robust multiarray analysis method (McCall *et al*, 2010). The differentially expressed genes were identified using the R package “limma”. Genes with *p* < 0.05 and a fold change of ≥ 1.5 (eATP synthase^high^ versus eATP synthase^low^) were considered differentially expressed genes. The signal-to-noise ratio was calculated to score the gene expression difference between cell lines with high and low abundance of eATP synthase, and the weighted Kolmogorov–Smirnov test was used to identify the significant gene sets.

### GO enrichment analysis and GSEA

We used GO-annotated gene sets to interpret the differentially expressed genes. The hypergeometric test was used to identify the over-represented GO terms, as follows:

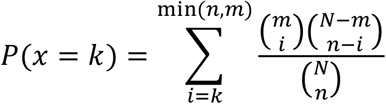

*N* is the number of genes in the genome, *m* is the number of genes in the genome that were annotated with a specific GO term, *n* is the number of differentially expressed genes, and *k* is the number of differentially expressed genes that were annotated with a specific GO term. The gene sets were curated from GO using Cytoscape software (Smoot *et al*, 2010). The gene list, pre-ranked according to fold change, was analyzed using GSEA (GSEA2-2.2.0) as previously described (Subramanian *et al*, 2005).

### Construction and purification of plasmids

The cDNA sequences encoding ATP5B or DRP1 were amplified by polymerase chain reaction (PCR; Thermal cycler modal: 2720; Applied Biosystems, Foster City, CA, USA) using the oligonucleotide primers (Genomics, Hsinchu, Taiwan) listed in Table S6. The amplified fragments were cloned into a paGFP-N1 vector (Addgene, Cambridge, MA, USA) or pCMV-HA-N vector (Clontech #635690), respectively. All clones were verified by sequencing (Genomics). The *Escherichia coli* DH5α strain was used as the competent cell for transformation of these constructs. The GeneJET Plasmid Midiprep Kit (Thermo Fisher Scientific) was used for the purification of plasmids, and the concentration of the purified plasmids was determined using the NanoDrop ND-1000 (NanoDrop Technologies, Montchanin, DE, USA). Plasmids were transfected into cells using jetPRIME (Polyplus-transfection, Illkirch, France) according to the instructions provided by the manufacturer, and the transfected cells were incubated for at least 24 h before further experiments.

### siRNA transfection

The siGENOME SMARTpool siRNAs were obtained from GE Healthcare (Dharmacon, GE Healthcare, Little Chalfont, Buckinghamshire, UK), and solubilized with nuclease-free water for further storage. Appropriate quantities of siRNAs were transfected into cells using commercial Lipofectamine 3000 (Invitrogen, Waltham, MA, USA) when the cells reached 60–70% confluency. In brief, the siRNAs and Lipofectamine 3000 reagent were diluted in Opti-Minimal Essential Medium (Thermo Fisher Scientific). The diluted siRNAs were mixed with the diluted Lipofectamine 3000 reagent and incubated at room temperature for 10–15 min. Finally, the siRNA–liposome mixture was added to the cells and the medium was replaced after 6 h of transfection to reduce toxicity.

### Western blotting

Proteins were denatured at 95°C for 5 min, separated by sodium dodecyl sulfate-polyacrylamide gel electrophoresis (SDS-PAGE) and transferred to a polyvinylidene difluoride membrane (PVDF, Millipore, Bedford, MA, USA). The PVDF membrane was blocked with 5% milk in Tris-buffered saline with Tween-20 (TBST) at room temperature for 1 h. After blocking, the proteins were hybridized with primary antibodies diluted in 5% milk/TBST at 4°C overnight. The membrane was further incubated with horseradish peroxidase-conjugated secondary antibodies (Abcam) diluted with 5% milk/TBST at room temperature for 1 h. The protein bands were analyzed using FluorChem M (ProteinSimple, San Jose, CA, USA).

### Flow cytometry

Cells were detached with 1 Mm EDTA (J.T. Baker, Center Valley, PA, USA) in PBS at room temperature for 5 min after incubation for at least 24 h. DMEM containing 10% fetal bovine serum was used for the inactivation of EDTA and resuspension of the cells. The resuspended cells were centrifuged at 300 × g at 4°C for 5 min and diluted to a concentration of 1 × 10^6^/ml using cold PBS. After fixing with 2% paraformaldehyde at 37°C in a water bath for 10 min, the non-permeable cells were incubated with the primary antibody against ATP synthase at 4°C overnight. Isotype IgG was loaded under the same conditions as a control. Further hybridization utilizing an Alexa488-conjugated goat anti-mouse or anti-rabbit IgG was performed at room temperature for 1 h. The labeled cells were washed with cold PBS and the signals were detected using a BD FACSCanto II instrument (BD Biosciences, San Jose, CA, USA).

### Real-time live imaging

Cells were seeded into 24-well plates and cultured for 48 h. The ATP5B-PaGFP plasmid was transfected using jetPRIME. Prior to the capture of live images, fluorescent probes were used to label the targeted organelles. Images were captured using a Zeiss LSM780 confocal microscope (Zeiss, Oberkochen, Germany) in an incubator at 37°C and 5% CO_2_. The paGFP signal was activated at a wavelength of 413 nm and detected at a 488 nm. Videos were captured every 4 s for 15 min to trace the movement of ATP synthase. Acquisition parameters were selected so as to allow the use of minimal laser intensity to prevent photobleaching. The movement distance of the ATP5B-paGFP signal was analyzed using Metamorph software (Molecular Devices, Sunnyvale, CA, USA). The colocalization coefficient between the ATP5B-paGFP signal and the PM was determined using Zeiss quantitative colocalization analysis software (Zeiss). The confocal images were displayed as a scatter plot.

### Immunofluorescence

Cells were seeded on the coverslips of a 12-well culture plate for 24 h. They were then fixed with 3.7% paraformaldehyde (Sigma–Aldrich) in PBS at room temperature for 20 min and washed thrice with PBS. To examine the expression of eATP synthase on cell surface, the cells were not permerabilzed. To detect the intracellular proteins, the cells were subsequently permeabilized using 0.1% triton X-100 (Sigma–Aldrich) in PBS. Next, the cells were incubated with 10% bovine serum albumin (BioShop) in PBS at room temperature for 1 h to block nonspecific binding. Cells were washed with PBS and hybridized with specific primary antibodies at 4°C overnight (16–18 h). After removing nonspecific binding by washing with PBS, the cells were incubated with Alexa488-conjugated goat anti-mouse or anti-rabbit IgG secondary antibodies at room temperature for 1 h. Finally, the coverslips were mounted onto the glass slides with 4’, 6-diamidino-2-phenylindole (DAPI) containing mounting medium (Thermo Fisher Scientific). The results were detected using fluorescence microscopy (Leica, Wetzlar, Hesse, Germany) and laser scanning confocal microscopy (Zeiss LSM 780 with an Airyscan detector and a Plan Apochromat 100 × /1.4 oil objective). The data were adjusted using the software Zen 2010 (Carl Zeiss Meditec AG, Jena, Thuringia, Germany).

### dSTORM imaging and analysis

Samples for super-resolution imaging were prepared according to the protocol of immunofluorescence. During imaging, samples were placed in an imaging chamber (Chamlide magnetic chamber, Live Cell Instrument, Seoul, Korea) and immersed in the imaging buffer containing 50 mM Tris-HCl and 10 mM NaCl (TN) buffer at pH 8.0 and an oxygen-scavenging system consisting of 60 mM mercaptoethylamine, pH 8.0, 0.5 mg/ml glucose oxidase, 40 μg/ml catalase, and 10% glucose (Sigma-Aldrich). The super-resolution imaging were performed on the direct stochastic optical reconstruction microscopy (dSTORM) system including a modified inverted microscope (Eclipse Ti-E, Nikon, Tokyo, Japan) with a 100 × oil immersion objective (1.49 NA, CFI Apo TIRF, Nikon) (Chong *et al*, 2020; Yang *et al*, 2018). Three light sources: a 637 nm laser (OBIS 637 LX 140 mW, Coherent, Santa Clara, CA, USA), a 561 nm laser (Jive 561 150 mW, Cobolt, Solna, Sweden), and a 405 nm laser (OBIS 405 LX 100 mW, Coherent) were homogenized (Borealis Conditioning Unit, Spectral Applied Research, Toronto, Canada) before focused onto the rear focal plane of the objective. For dual-color imaging, Alexa Fluor 647 and Cy3B were excited by a 647 nm and a 561 nm laser lines, respectively, operated at 3−5 kW/cm^2^. A 405 nm light was used to convert a small fraction of the fluorophores from a quenched state to an excited state. The single-molecule signals were registered on an electron-multiplying charge-coupled device (EMCCD) camera (Evolve 512 Delta, Photometrics, Tucson, AZ, USA). 10,000–20,000 frames were acquired for an exposure time of 20 ms. Each single-molecule burst was localized using a MetaMorph Super-resolution Module (Molecular Devices, Sunnyvale, CA, USA) and cleaned with a Gaussian filter with a radius of 1 pixel.

### Mitochondrial image analysis

We analyzed the captured images using Icy to quantify the perimeter and area of mitochondria (deChaumont et al, 2012). Firstly, we loaded the images into Icy (http://icy.bioimageanalysis.org/), and the weak signals were adjusted to enhance image contrast. Subsequently, images were blurred for the networked mitochondria using a Gaussian filter tool, and displayed to check the results. The filtered results were loaded to the K means thresholder for automatic adjustment, and the binary images were output by inputting the images to the thresholder and label extractor (Pagliuso et al., 2016). Next, the processed images were loaded into a spot detector with parameter setting to three-pixel detection. The results revealed the perimeters, areas, and contour level of every detected spot (DeVos & Sheetz, 2007; Mitra & Lippincott-Schwartz, 2010).

### Statistical analysis

All data were presented as the mean ± SD of three experimental replicates and analyzed using Student’s unpaired two-tailed *t*-test. A *p*-value of < 0.05 denoted statistically significant differences.

## Supporting information

Extended View Figures

## Data availability

All original mass spectrometry data have been deposited to the ProteomeXchange Consortium via the PRIDE partner repository (Vizcaíno *et al*, 2016) with the dataset identifiers PXD006791 (plasma membrane proteome), PXD007036 (mitochondrial proteome), and PXD010784 (KIF5B-co-immunoprecipitation proteome).

## Acknowledgments

We thank Technology Commons in College of Life Science, National Taiwan University for technical assistance. We thank Prof. David Chan (Division of Biology and Biological Engineering, California Institute of Technology) for the constructive suggestions. We would like also to thank Amber Kao and Anthony Abram for editing and proofreading this manuscript. This work was supported by the Ministry of Science and Technology (MOST 105-2320-B-002-057-MY3, MOST 106-2320-B-002-053-MY3, MOST 107-2321-B-006-020, MOST 107-2221-E-010-017-MY2, MOST 109-2221-E-010-012-MY3, MOST 109-2320-B-002-017-MY3), the Higher Education Sprout Project (NTU-108L8807A and 109L8837A) and the National Health Research Institutes (NHRI-EX109-10709BI) in Taiwan.

## Author Contributions

Y.-W. C., H.-C. H., and H.-F. J. conceived this study. Y.-W. C. and T. T. Y. designed and performed the super-resolution imaging. Y.-W. C., M.-C. C., and T.-Y. H. designed and performed all immunofluorescence and flow cytometry experiments. Y.-W. C. and Y. L. designed and performed the real-time live-cell imaging. C.-F. Y. designed and performed all the cloning in this study. Y.-W. C. and J.-T. H. performed the homology modeling and protein-protein docking simulation. Y.-H. H. and C.-L. H. performed computational data analyses. Y.-W. C. and H.-F. J. analyzed and interpreted the results. Y.-W. C., T. T. Y., H.-C. H., and H.-F. J. wrote and proofread the manuscript.

