## Extended View Figures for "Mitochondrial ATP Synthase Trafficking along Microtubules to Cell Surface Depends on KIF5B and DRP1"

**Figure EV1.**

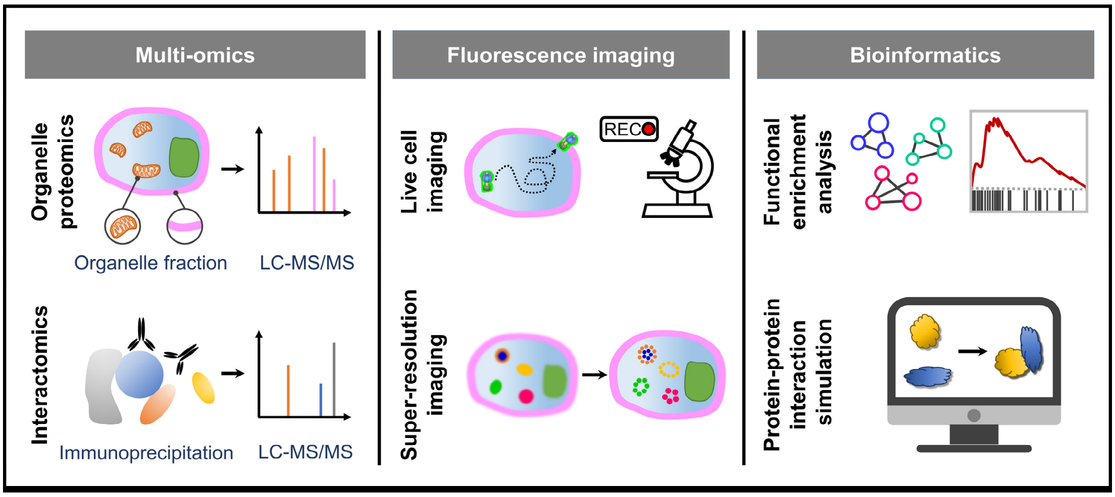


**Figure EV1. Schematic of experiment design for identifying the trafficking mechanism of mitochondrial ATP synthase.**

#### Figure EV2.

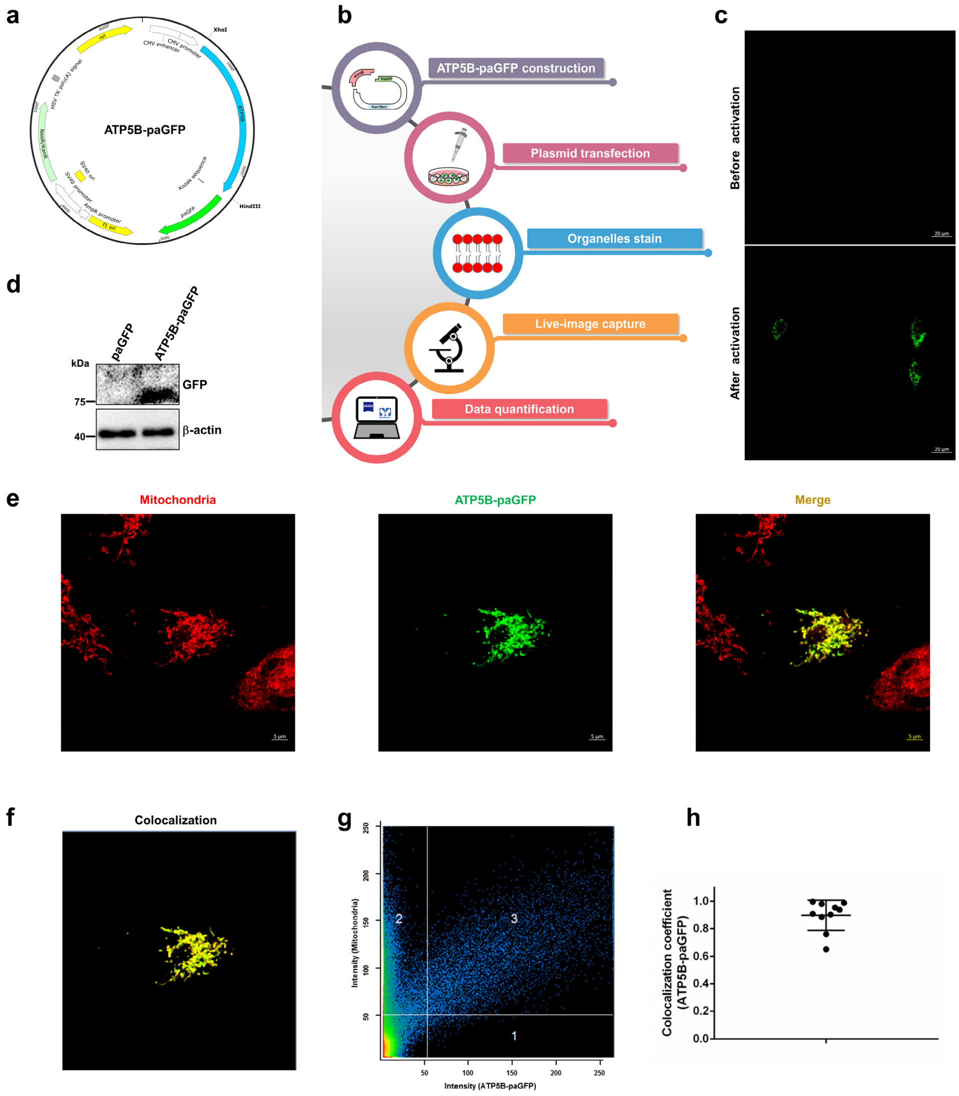


**Figure EV2. (related to Fig1): The spatial localization of ATP synthase in live A549 cells is determined using the ATP5B-paGFP fusion protein.**

A The sequence map for ATP5B-paGFP recombinant construct.

B Schematic representation of the real-time tracing system of ATP synthase.

C The ATP synthase β-paGFP signal in live cells was photoactivated with a 405 nm laser using confocal microscopy. Scale bars, 20 µm.

D The expression levels of ATP synthase β-paGFP fusion proteins in transfected cells were detected by western blotting using antibody against GFP.

E The subcellular localizations of intracellular ATP synthase β-paGFP fusion protein (green) and the mitochondria (MitoTracker; red) in live cells were detected using confocal microscopy. The colocalization of these two signals is shown as yellow signals in the merged images. Scale bars, 5 μm.

F The merged images in E were further processed using ZEN software to enhance image contrast.

G The fluorescent signals of ATP synthase β-paGFP fusion protein (green; area 1), mitochondria (red; area 2), and colocalization (yellow; area 3) were determined by ZEN software and are shown as the scatter plot.

H The colocalization coefficient of ATP synthase β subunit-paGFP and mitochondria (colocalization fluorescent area versus total ATP synthase β subunit-paGFP fluorescent area) was presented as mean ± SD in bar chart (n = 10). * $p < 0.05$, ** $p < 0.01$, *** $p < 0.001$.

#### Figure EV3.

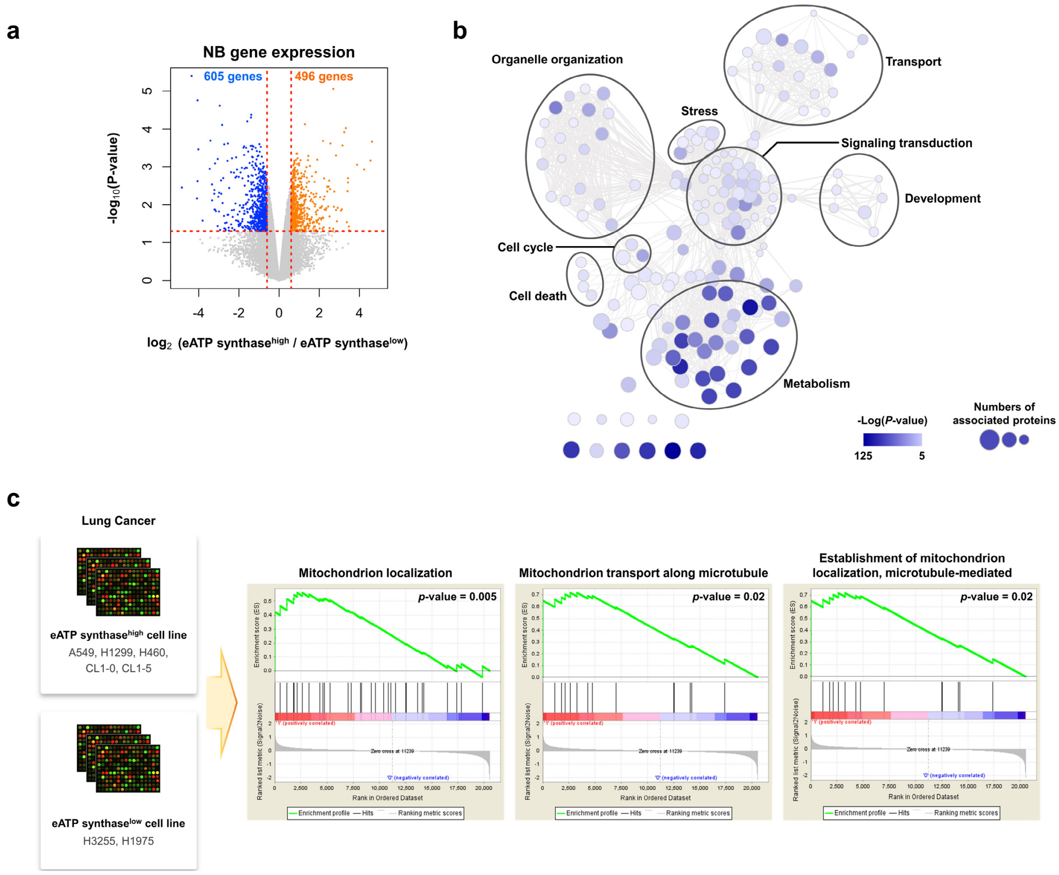


**Figure EV3. (related to Fig 2): The potential mechanism of ATP synthase trafficking to the cell surface.**

A Volcano plot depict the gene expression differences between eATP synthase$^{high}$ and eATP synthase$^{low}$ cell lines. The orange points represent the significantly up-regulated genes and the blue points represent the significantly down-regulated genes ($p < 0.05$ and fold-change > 1.5).

B Functional enrichment network of the eATP synthase-related genes. Nodes represent gene ontology terms (GO terms) which were statistically enriched in the genes with differential expression. Links connecting nodes indicate the relatedness between nodes.

C Gene set enrichment analysis (GSEA) was conducted on the gene expression profiles of the eATP synthase$^{high}$ and eATP synthase$^{low}$ lung cancer cell lines. The gene sets associated with mitochondrial trafficking were significantly positively enriched (*p*-values < 0.05). Enrichment scores are 0.72, 0.72, 0.56 for GO terms “Mitochondrion transport along microtubule”, “Establishment of mitochondrion localization, microtubule-mediated” and “Mitochondrion localization”.

### Figure EV4.

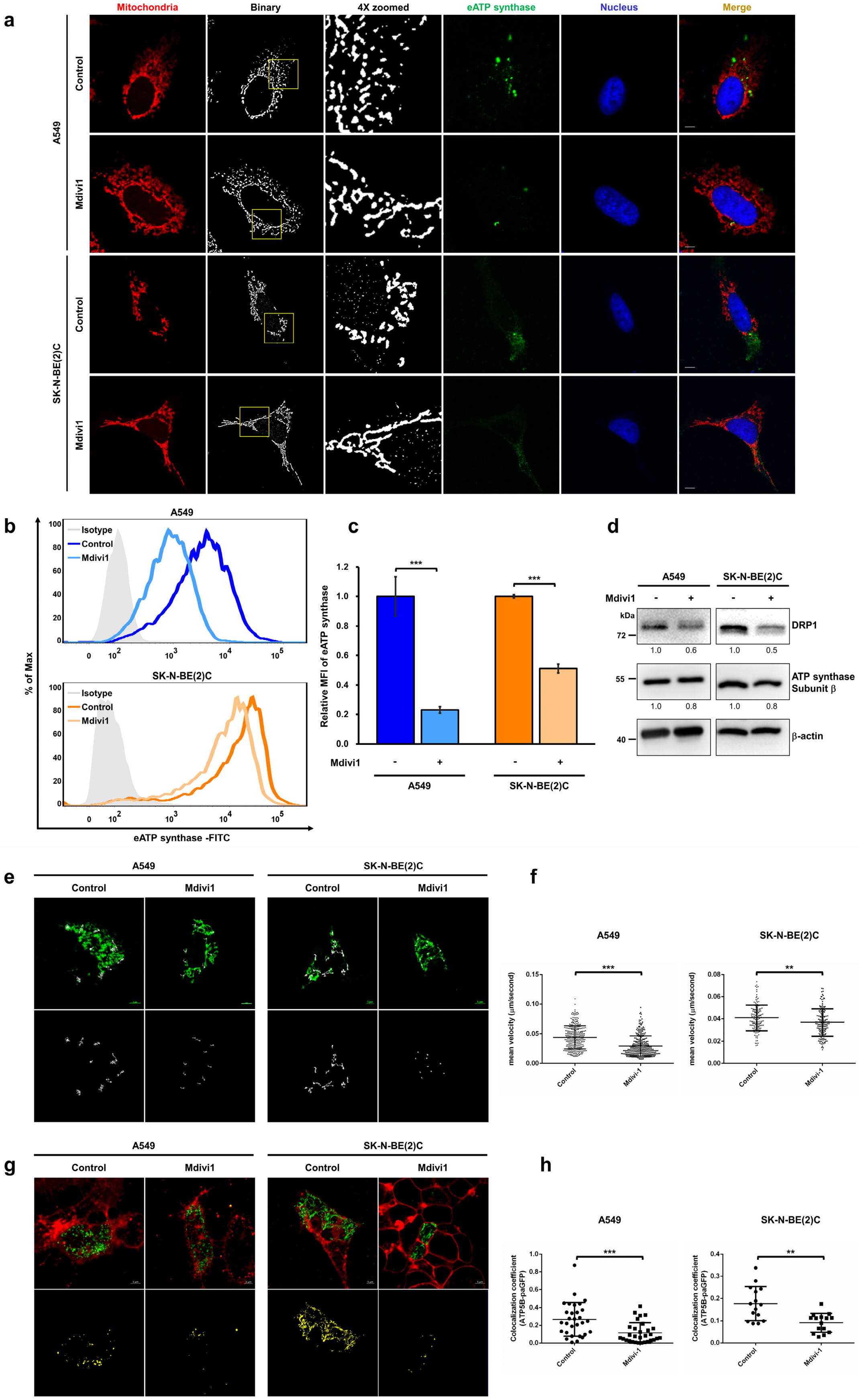

a
Mitochondria
Binary
4X zoomed
eATP synthase
Nucleus
Merge
A549
Control
Mdivi1
SK-N-BE(2)C
Control
Mdivi1
b
A549
Isotype
Control
Mdivi1
SK-N-BE(2)C
% of Max
eATP synthase -FITC
c
Relative MFI of eATP synthase
***
Mdivi1
A549
SK-N-BE(2)C
d
A549
SK-N-BE(2)C
Mdivi1
kDa
DRP1
ATP synthase Subunit β
β-actin
1.0 0.6 1.0 0.5
1.0 0.8 1.0 0.8
e
A549
SK-N-BE(2)C
Control
Mdivi1
f
A549
SK-N-BE(2)C
mean velocity (μm/second)
Control
Mdivi-1
g
A549
SK-N-BE(2)C
Control
Mdivi1
h
A549
SK-N-BE(2)C
Colocalization coefficient (ATP5B-paGFP)
Control
Mdivi-1

**Figure EV4. (related to Fig 4): Inhibition of mitochondrial fission suppresses the transport of ATP synthase toward the cell surface.**

A A549 and SK-N-BE(2)C cells were treated with DMSO or Mdivi1 (30 μM). Mitochondrial morphology and the expression of eATP synthase on the cell surface of A549 and SK-N-BE(2)C cells were visualized using confocal microscopy (100×) and Mitotracker (red) or the antibody against the ATP synthase complex, followed by hybridization with Alexa 488 anti-mouse IgG (green). The binary images shown were processed using Icy software. Scale bars, 5 μm.

B The abundance of eATP synthase was determined via flow cytometry using anti-ATP synthase antibody in nonpermeable cells.

C The relative MFI of eATP synthase based on the flow cytometry data is displayed as a bar chart. Data presented are the mean ± SD (n = 3).

D The protein levels of Drp1 and ATP synthase subunit β were verified using western blotting in the DMSO- and Mdivi1-treated groups. The intensity of the bands was normalized to that of β-actin and relative to the expression of each protein in the DMSO-treated group.

E The movement of the ATP synthase subunit β-paGFP fusion protein in the DMSO or Mdivi1 group was recorded for 15 min via confocal microscopy (upper). Their tracks are shown as white lines and were analyzed using Metamorph software (lower).

F The dot plot represents the quantification of the translocation velocity of both A549 and SK-N-BE(2)C cells (n = 30).

G The localization of ATP synthase subunit β-paGFP fusion protein (green) and PM (CellMask; red) were determined using confocal microscopy. The colocalization of these two signals is shown as yellow fluorescence in the merged images (upper), and was further processed using ZEN software (lower).

H The dot plot shows the colocalization coefficient (colocalization fluorescent area versus total ATP synthase subunit β-paGFP fluorescent area) ($n = 30$). Values shown are the mean ± SD. * $p < 0.05$, ** $p < 0.01$, *** $p < 0.001$
